# The structure of the *Physcomitrium Patens* Photosystem I Reveals a Unique Lhca2 Paralogue replacing Lhca4

**DOI:** 10.1101/2021.11.26.470156

**Authors:** C Gorski, R Riddle, H Toporik, Z Da, Z Dobson, D Williams, Y Mazor

**Author notes:** Equally contributed (C Gorski, R Riddle, H Toporik).

## Abstract

The moss *Physcomitrium patens* diverged from green algae shortly after the colonization of land by ancient plants. This colonization posed new environmental challenges which drove evolutionary processes. The photosynthetic machinery of modern flowering plants is adapted to the high light conditions on land. Red shifted Lhca4 antennae are present in the photosystem I light harvesting complex of many green lineage plants but absent from *P. patens*. The Cryo-EM structure of the *P. patens* photosystem I light harvesting complex I supercomplex (PSI-LHCI) at 2.8 Å reveals that Lhca4 is replaced by a unique Lhca2 paralogue in moss. This PSI-LHCI supercomplex also retains the PsaM subunit, present in cyanobacteria and several algal species but lost in higher plants, and the PsaO subunit responsible for binding light harvesting complex II. The blue shifted Lhca2 paralogue and chlorophyll *b* enrichment relative to higher plants make the *P. patens* PSI-LHCI spectroscopically unique among other green lineage supercomplexes. Overall, the structure represents an evolutionary intermediate PSI with the crescent shaped LHCI common in higher plants and contains a unique Lhca2 paralogue which facilitates the mosses adaptation to low light niches.

## Introduction

Oxygenic photosynthesis which produces the oxygen in the atmosphere and the energy for CO_2_ fixation evolved in cyanobacteria more than 2.5 billion years ago^1,2^. Approximately 1.5 billion years later, the primary endosymbiotic event establishing the green lineage took place as the non-photosynthetic ancestor of modern green algae engulfed a cyanobacterium^3^. Following this primary endosymbiotic event, ancient bryophytes, the ancestors of the moss *Physcomitrium patens* (*P. patens*, also known as *Physcomitrella patens*), colonized land from the sea and established the green lineage as the primary land dwelling photosynthetic autotrophs^4–6^ leading to modern flowering plants, known as angiosperms, the youngest members of the green lineage.

The transition to land posed new challenges for early plants. For sessile organisms, light on land was intense and inescapable, and water was no longer ubiquitous. In order to deal with constant solar radiation early plants developed enhanced DNA repair mechanisms, desiccation tolerance, and an increased repertoire of chlorophyll binding proteins^4^. These changes in light conditions also necessitated photosynthetic adaptations.

The photosynthetic machinery, responsible for light absorption and its conversion to useable energy utilized in carbon fixation, includes two photosystems, photosystem I (PSI) and photosystem II (PSII). PSI and PSII are pigment-protein complexes that absorb light and utilize its energy to facilitate electron transfer reactions. PSII oxidizes water and the extracted electrons are transferred through additional components of the photosynthetic electron transport chain to PSI which catalyzes electron transfer from the electron donor, plastocyanin to ferredoxin. This process produces the reducing power and the energy needed for carbohydrate assimilation^7–9^.

In the green lineage, PSI is found as a monomer and consists of a remarkably conserved core bound to light harvesting complexes (Lhc) forming the PSI-LHCI complex. LHCI functions as antenna to increase the amount of absorbed photons^10–14^. The Lhc proteins containing three transmembrane helices with a highly conserved core, are encoded by a large multigene family and interact with PSI (Lhca) and PSII (Lhcb)^15–17^.

In angiosperms, six Lhca (1-6) gene types can be found. The PSI-LHCI complex of the angiosperms *Pisum sativum* and maize have been shown to consist of four LHC proteins made up from two heterodimers, Lhca1/Lhca4 and Lhca2/Lhca3^12,13,18-21^. Lhca5 and Lhca6 are present at sub stoichiometric amounts and increase in abundance under high light conditions^15,22^. Lhca5 and Lhca6 play a role in the formation of the PSI-NDH supercomplex associated with cyclic electron flow^23–25^. It was shown in *Arabidopsis thaliana* (*A. thaliana*), that different light regimes did not change the stoichiometry of the LHC antenna^26^ and that locations of specific Lhca proteins in plants are not interchangeable, however Lhca5 can replace Lhca4 in Lhca4 knockout plants^27^.

In the single celled green alga *Chlamydomonas reinhardtii* (*C. reinhardtii*) nine Lhca genes (Lhca1-9) are expressed and are part of PSI-LHCI super-complex^28–30^. There is a high degree of diversity in the structures of PSI-LHCI complexes from green algae, ranging from four and six LHC proteins in *Dunaliella salina*^31,32^, to ten LHC proteins in *C. reinhardtii*^33,34^ and the macroscopic green algae *Bryopsis corticulans*^35^.

Interestingly, the genes for Lhc proteins in the *P. patens* genome are more diverse and redundant in comparison to green algae and higher plants, with several paralogous genes for Lhca1, Lhca2 and Lhca3. This is partly explained by a genome duplication event which occurred in the moss 45 million years ago^36,37^. The moss Lhca1, Lhca2, Lhca3 and Lhca5 are similar to their plant counterparts^38,39^. However, Lhca4 and Lhca6 are missing in the *P. patens* genome^5^.

EM analysis at low resolution of PSI complexes isolated from *P. patens* suggests two different PSI-LHCI complexes; one that resembles the plant structure and includes four Lhc proteins, and a larger alga-like complex^36,40,41^. The larger complex contains LHCII - a PSII antenna, two layers of LHCI belts, where the outer belt is shifted in comparison to the alga complex and Lhcb9, a moss specific Lhc protein^40,41^. Lhcb9 knockout mutants impair the formation of this larger complex^37,42^.

A unique feature of the LHCI complex is the presence of low energy chlorophylls among the antenna chlorophylls (Chls), also known as red Chls. These Chls absorb light in longer wavelengths than 700 nm, resulting in a red shifted absorption spectrum^43,44^. Red Chls were suggested to play a role in light harvesting by extending PSI absorption as well as playing a role in photoprotection^45–47^. *In vitro* reconstitution of Lhca proteins has characterized the red Chl content and spectral properties of each Lhca protein. The low temperature emission spectrum peaks of each Lhca protein are 690nm, 702nm, 725nm and 733nm for Lhca1-4, respectively^21,48,49^. The heterodimers in the plant LHCI belt are “blue” and “red” partners, where Lhca1 and Lhca4 form one dimer and Lhca2 and Lhca3 form another^50^. In contrast, *P. patens* does not code for Lhca4, the lowest energy form of the Lhca proteins in the plant antenna. The composition of the moss LHCI complex without Lhca4 remains to be determined.

Here we report the Cryo-EM structure of the moss *P. patens* PSI-LHCI complex with four Lhca proteins. The structure determines the identity of the Lhc protein that occupies the Lhca4 position as an Lhca2 paralogues. The participation of this paralogue in the moss LHCI and the amount of Chl *b* incorporated into the antenna are the underlying reasons for the spectroscopic differences observed between the higher plant and the moss PSI-LHCI. We suggest that this structure represents the adaptation of the photosynthetic machinery to the current low light habitat of mosses.

## Results

### Overall PSI-LHCI structure

We solved the structure of the PSI-LHCI complex from the moss *P. patens* using single particle Cryo-EM. The density map of the major class was reconstructed from 114,307 particles to 2.8 Å resolution allowing precise assignment of 17 subunits, 159 chlorophyll molecules, 35 carotenoids and 12 lipids (**figure 1a** and **supplementary figures 1, 2** and **supplementary table 1**). Overall, the structure of the PSI-LHCI complex is similar to the pea PSI-LHCI complex with a monomeric core complex including twelve subunits, with two modifications. PSI from *P. patens* still carries the PsaM subunit which is not found in higher plants. PsaM is not found in the single celled green alga *C. reinhardtii*^33,34^ and *Dunalliela Salina*^32^ but is present in the macroscopic green alga, *Bryopsis corticulans*^35^ and the red alga, *Cyanidioschyzon merolae*^51,52^, suggesting that it was lost multiple times during the evolution of green lineage PSI and the moss PSI-LHCI is the last member of the multicellular lineage leading to higher plants that carry PsaM.

**Figure 1.**
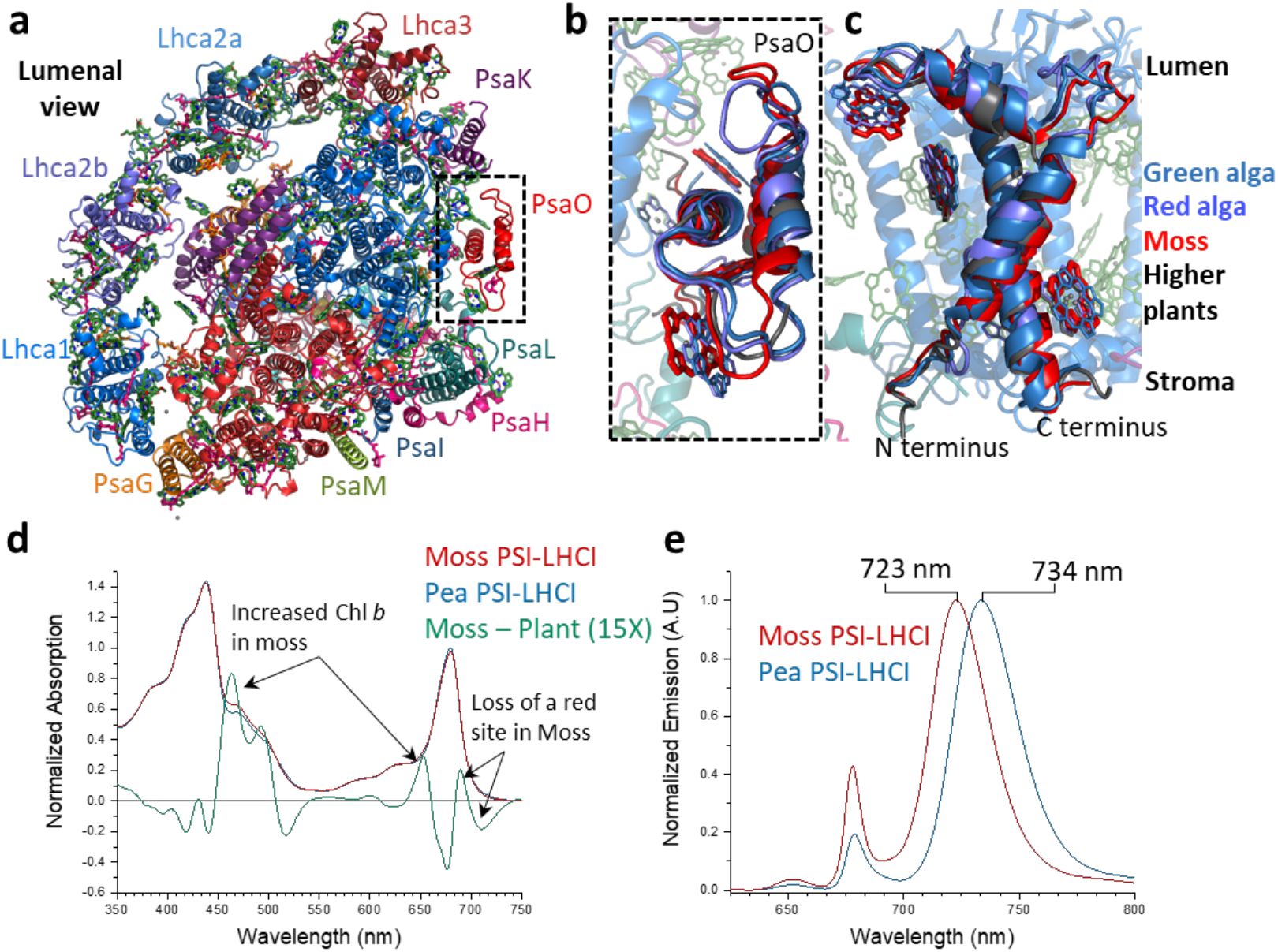
The overall structure of moss PSI-LHCI complex. **a.** A view from the lumen of PSI-LHCI from *P. patens.* Two forms of Lhca2 are present in LHCI. **b** and **c**. Luminal and side views comparing PsaO across the green lineage, from higher plants (PDB ID 5ZJI), green alga (PDB ID 6SL5), red alga (PDB ID 5ZGB) and moss (PDB ID 7KSQ). The comparison shows large variations in Chl coordinating loops and a conserved core structure. **d**. Absorption spectra of pea (blue) and moss (red) PSI-LHCI with the difference spectra shown in green. Increased Chl *b* content and decreased far red absorption are clearly seen. **e**. 77 K emission also indicates the loss of a red site from LHCI in the moss compared to higher plants.

Examination of the overall map at the region occupied by PsaO in higher plants revealed very weak density in its position. Focused classification of the particle set on this region resulted in a subclass of particles (approximately 13%, **supplementary figures 2** and **3**) that contained PsaO together with an additional density in its proximity, this map region was refined to a resolution of 3.8 Å (**supplementary figure 3**). This additional density is best fitted as an unidentified Lhc monomer with two modifications. First, the N terminus is not resolved and is probably degraded, second, the membrane orientation is opposite to the native orientation, with the N and C terminal ends located at the luminal and stromal sides of the membrane (inverted compared to other Lhc’s). Together, this suggests that the association between this antenna and PSI-LHCI at this site occurs post solubilization and it was not investigated further (**supplementary figure 3**). PsaO was previously undetected in the moss PSI-LHCI complex using antibodies against the plant PsaO^36^. The co-occurrence of the inverted Lhc and PsaO suggests that in the moss, and possibly in higher plants, PsaO is a very general adaptor protein to PSI. Comparing PsaO across the green lineage shows high structural conservation in the membrane region, with high variability in the two luminal loops of the subunit (**figure 1b and c**). In addition, another subpopulation of the particles appeared as LHCI devoid of Lhca3 (**supplementary figure 2b**). It is presently not clear if this form of PSI-LHCI represents a *bona fide* cellular form of PSI-LHCI or results from the isolation procedure.

A major trend in the evolution of PSI-LHCI in the green lineage is the appearance of low energy or red Chls in LHCI. The moss PSI-LHCI is a very clear intermediate of this process^5^. Comparing the absorption spectra of PSI-LHCI purified from moss and higher plants (*Pisum sativum*, pea) shows that both complexes are similar and points to two clear differences (**figure 1d**). First, in the red region of the Q_y_ Chl *a* transition, the shape of the difference spectra shows a clear minimum compared to higher plant PSI-LHCI, suggesting that some red sites are missing (**figure 1d**). Second, the positive peaks around 650 nm and 460 nm suggests that the Chl *b* content of the moss PSI-LHCI is higher than that of the higher plant complex (**figure 1d**).

In plants, Lhca4 was found to contain the lowest energy form of Chl *a* in LHCI. The 77 K emission spectra of the moss PSI-LHCI is markedly blue shifted compared to that of higher plants and this suggests that at least one of the red Chls sites in LHCI is missing. *P. patens* does not contain any genes for Lhca4 and it was previously suggested that *P. patens* PSI interacts with a smaller antenna ^5^ or that another Lhca protein occupies its position in LHCI^36^.

### The composition of the moss LHCI antenna

The genome of *P. patens* contains four Lhca families (Lhca1, 2, 3 and 5) with a large number of paralogs for each making the assignment of antenna subunits challenging^5^. In *Arabidopsis thaliana*, deletion of the Lhca4 gene leads to the binding of Lhca5 to LHCI^27^. In *P. patens*, semi quantitative proteomic data of the PSI-LHCI complex showed that Lhca5 is expressed in sub stoichiometric amounts leading to the conclusion that different isoforms of Lhca1, Lhca2 or Lhca3 potentially occupy the Lhca4 location^36^. To identify the Lhca occupying this position we constructed a phylogenetic tree from all annotated Lhca genes from a selection of green lineage genomes and structures (**supplementary figure 4a**) including the angiosperms *P. Sativum, Zea mays, A. thaliana*, the lycophyte *Selaginella moellendorffii*, the bryophytes *P. patens, Marchantia polymorpha, Sphagnum fallax*, and the chlorophytes *C. reinhardtii, B. corticulans* and *D. Salina*. This analysis identified three Lhca1 genes, four Lhca3 genes, one Lhca5 gene, and seven Lhca2 genes in *P. patens*. None of these paralogs clustered with the higher plant Lhca4 genes, as expected. The seven Lhca2 genes clearly belonged to two different classes which we named Lhca2a and Lhca2b (**supplementary figure 4b and c**).

During the assignment process all isoforms of Lhca1, Lhca3 and Lhca5 were clearly ruled out due to the presence of unique loops absent from the density map, leaving the two Lhca2s as the final candidates to occupy the plant Lhca4 position in the moss LHCI. **figure 2** shows the alignment of Lhca2a vs Lhca2b genes from *P. patens*. All 45 sequence positions that distinguish these two families and are present in the final model are marked. The majority of these positions (68%, colored in green in **figure 2**) support the assignment, while 14 positions (32%) cannot be distinguished by the map alone and are neutral. No positions that support the opposite assignment were observed and map sections of the clearest examples are shown next to the alignment (**figure 2**). We conclude that each Lhca2 family occupies a distinct position in LHCI; Lhca2a in the plant Lhca2 position and Lhca2b in the plant Lhca4 position.

**Figure 2.**
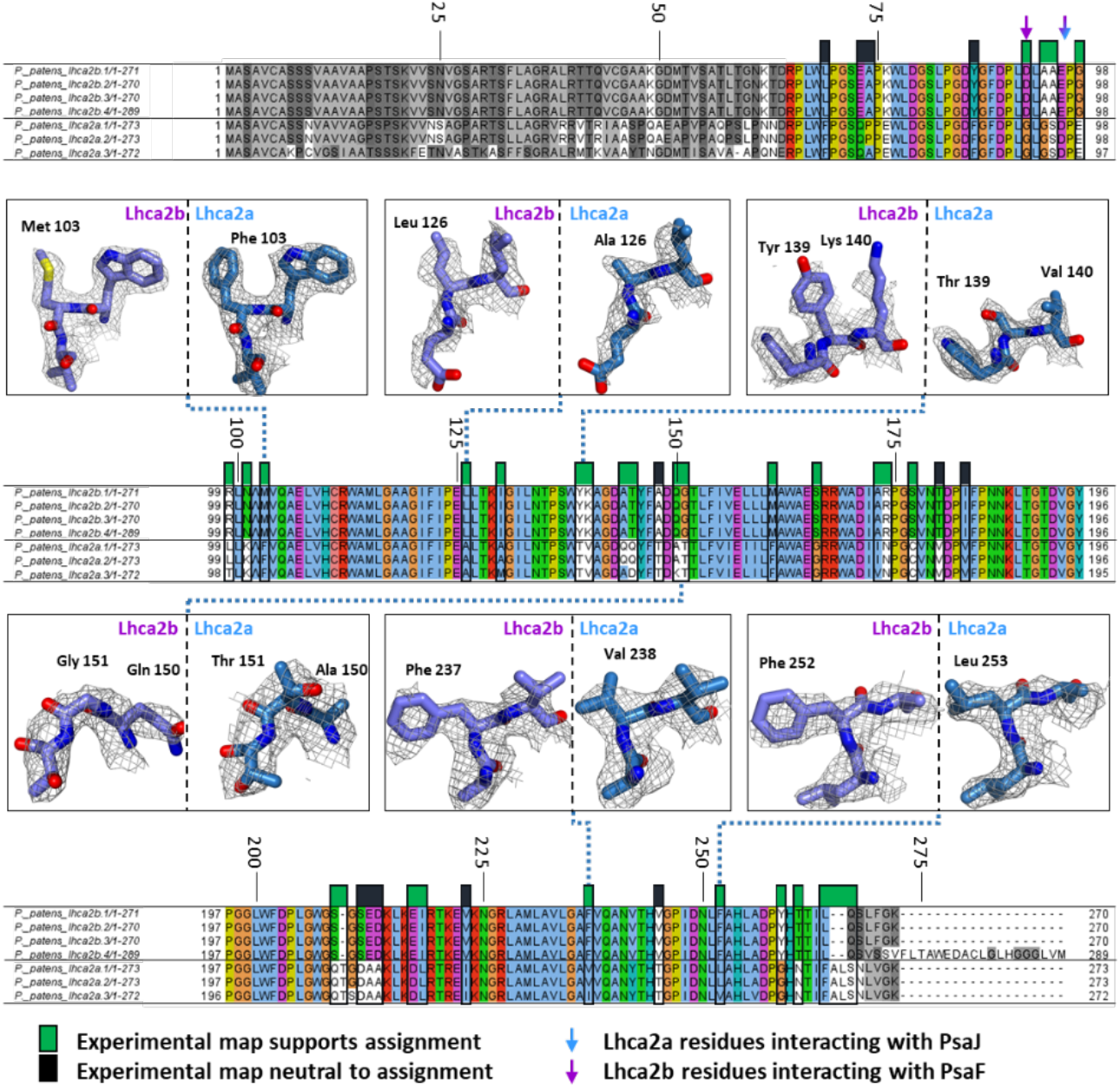
The *P. patens* Lhca2b occupies the plant Lhca4 position in the LHCI antenna. The alignment of all seven Lhca2 genes is shown with all amino acid differences in boxes. The gray parts of the alignment are not modeled in our final structure. The boxes labeled in green show well-defined densities supporting the assignment of each Lhca2 isoform in its position. The positions neutral to the assignment are labeled in black. Selected examples of differentiating residue fits are shown.

### Site specificity determinants of Lhca2a and Lhca2b - co-evolution of antenna and peripheral PSI subunits to achieve specific binding

Experimental data has shown that Lhca’s are not interchangeable^27^. What are the elements that drive the specific binding of individual Lhca’s to their position in LHCI? The PsaF side of PSI binds the LHCI antenna, PsaB, PsaF, PsaJ, PsaA and Psak participate in the interactions with Lhca1, Lhca2b, Lhca2a and Lhca3, respectively. The interacting residues between Lhca1 and PsaB or Lhca3 and PsaA are conserved between higher plants and moss and appear in their final form in the moss. The critical interactions between Lhca2a and Lhca2b and PSI occur at the stromal side at two small interaction surfaces with PsaJ and PsaF respectively (**figure 3a**).

**Figure 3:**
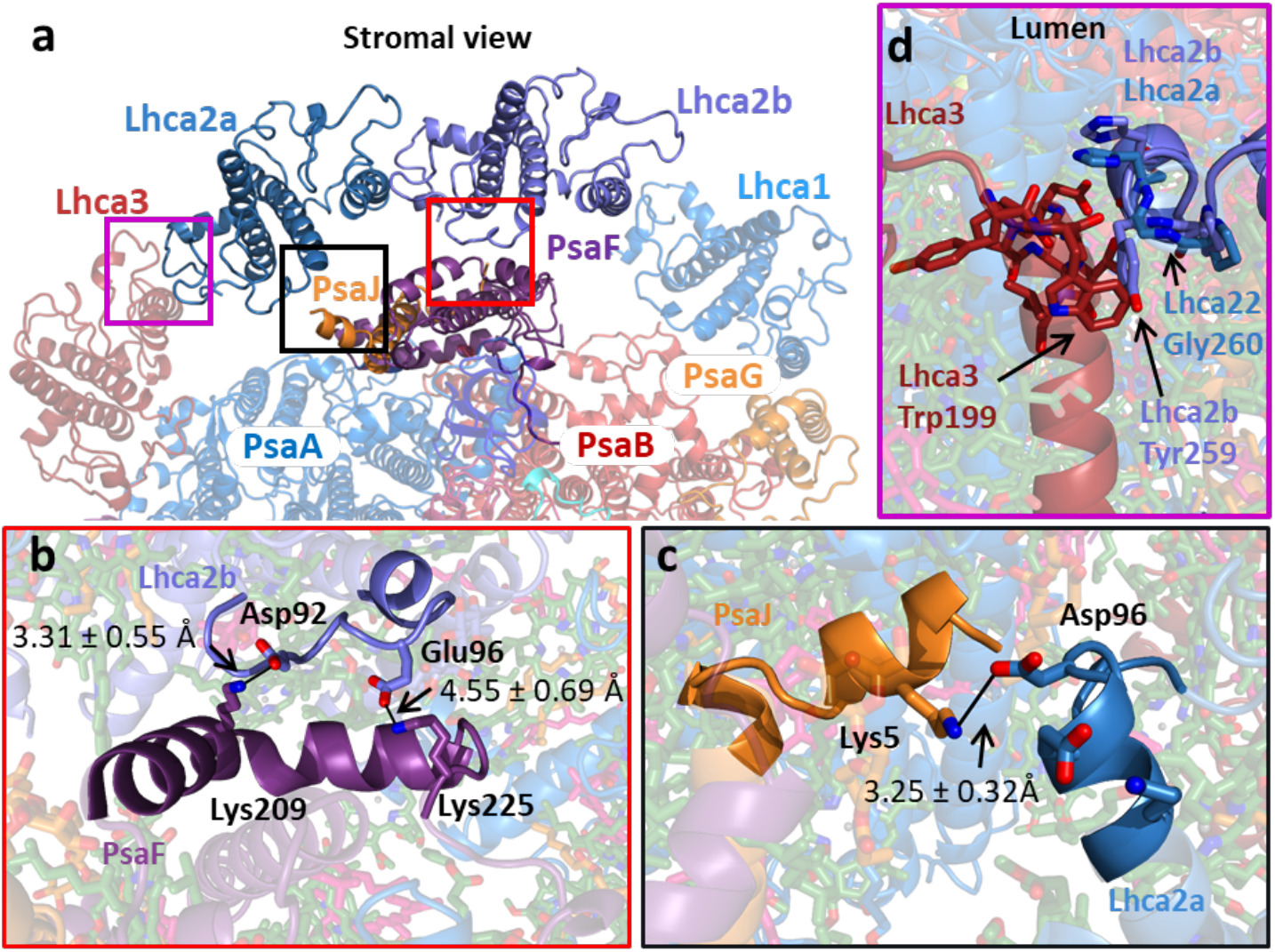
Antennae specificity in LHCI. **a.** Stromal view of the PSI-LHCI interface. The Lhca3-Lhca2a interface is boxed in magenta, the Lhca2a-PsaJ Interface is boxed in black and the Lhca2b-PsaF interface is in red. **b**. Two salt bridges are responsible for the interaction between Lhca2b (light purple) and PsaF (dark purple); Lhca2b Asp92 with PsaF Lys209 and Lhca2b Glu96 with PsaF Lys225. **c**. At the Lhca2a PsaJ Interface, Lhca2a (blue) Asp96 forms a salt bridge with PsaJ (orange) Lys5. **d.** Superposition of Lhca2b (light purple) on Lhca2a (blue) location emphasizes the steric clash between Lhca4 Tyr259 and Lhca3 (red) Trp199 preventing Lhca2b from occupying the Lhca2a position. All distances between residues were weighted based on rotamer probabilities (see method section for details).

The mode of interaction is the same, both interfaces form a salt bridge between positively charged residues on PSI and negatively charged residues in the respective antennae. Both Lhca2a and Lhca2b contain a negatively charged residue in position 96 which completes a single salt bridge (**figure 3b** and **c**). This interaction may be responsible for the initial binding predating the evolution of Lhca2b. In addition to this bridge, Lhca2b contains an additional negatively charged residue which makes up a second salt bridge with a lysine residue on PsaF (**figure 3b**). Interestingly, this interaction is present in the plant Lhca4 as well. The higher plant PsaF contains an arginine residue at the equivalent position preserving the positive charge. However, the plant Lhca4 completes the salt bridge using a glutamate residue located four amino acids downstream^12^. This glutamate residue is replaced by an alanine in the moss Lhca2b (**figure 2-residue 95**). This general trend of positively charged residues on PSI forming electrostatic interactions with negatively charged residues on Lhca’s repeats itself for Lhca3 as well. Lumenal interactions between individual Lhca’s contribute to specific binding of Lhca2b. These interactions are formed between the C-terminus of Lhca2b and a small patch at the beginning of the second trans-membrane helix of Lhca2a. Here we mainly find that Lhca2b cannot bind the Lhca2a positions due to the presence of a tyrosine residue present in Lhca2b (**figure 3d**).

To summarize, the highly conserved core subunits PsaA and PsaB maintain a specific set of interactions with the antennae Lhca1 and Lhca3, while the peripheral subunits PsaF and PsaJ determine the isoforms of Lhca that will interact with PSI. Only one isoform exists for the chloroplast encoded subunits PsaA, PsaB and PsaJ in the moss. However, four isoforms can be found for the nuclear encoded subunit, PsaF. Nevertheless, the sequence of the interaction sites is the same in all isoforms and determines the identity of the specific Lhca at this location.

### Evolution of the red antenna state along the green lineage

The location of red Chls across PSI is species dependent and can be found in the PSI core and in the LHCI complex. In the plant model organism, *A. thaliana*, the red Chls are located in the LHCI complex. Mutagenesis studies and biochemical reconstructions have characterized the spectroscopic properties of each Lhca protein and identified the specific pair of Chls that contribute to the red shifted absorption^49,53,54^. Lhca3 and Lhca4 carry the lowest energy Chls in the antenna with characteristic emission peaks at 725nm and 735nm, respectively^50^. These red shifted characteristics originate from excitonic interactions between two Chls; Chl 603 and Chl 609. An asparagine residue coordinating Chl 603 is crucial for the apparent red shifted absorption. Replacing this residue with a histidine residue led to a reduced far-red absorption. In addition, Chl 603 is coordinated by a histidine residue in Lhca1 and Lhca2 which do not contain red shifted Chls^53^.

The moss PSI-LHCI structure shows that Lhca3 Chl 603, coordinated by asparagine, and its immediate chemical environment is virtually identical between the moss and the higher plant complexes preserving the far-red emission characteristics. (**figure 4a**). In contrast, Chl 603 in Lhca2b is coordinated by a histidine residue, while in Lhca4 this Chl is coordinated by asparagine forming the lowest energy form of Chl *a* in the higher plant antenna. In addition, replacement of a phenylalanine residue with a methionine results in a small (0.78 Å displacement) shift of the Chl ring which contributes to the modification of the spectroscopic properties of this Chl dimer (**figure 4b**). In the plant Lhca’s, an additional gap Chl (Chl 617) is coordinated next to both the Lhca3 and Lhca4 red dimers^20,55^. Despite the fact that Lhca2b occupies the same position as Lhca4, we find that this gap Chl is missing, in correlation with the absence of a low energy site, suggesting that the function of Chl 617 may be related to the red state of Lhca4 and Lhca3.

**Figure 4.**
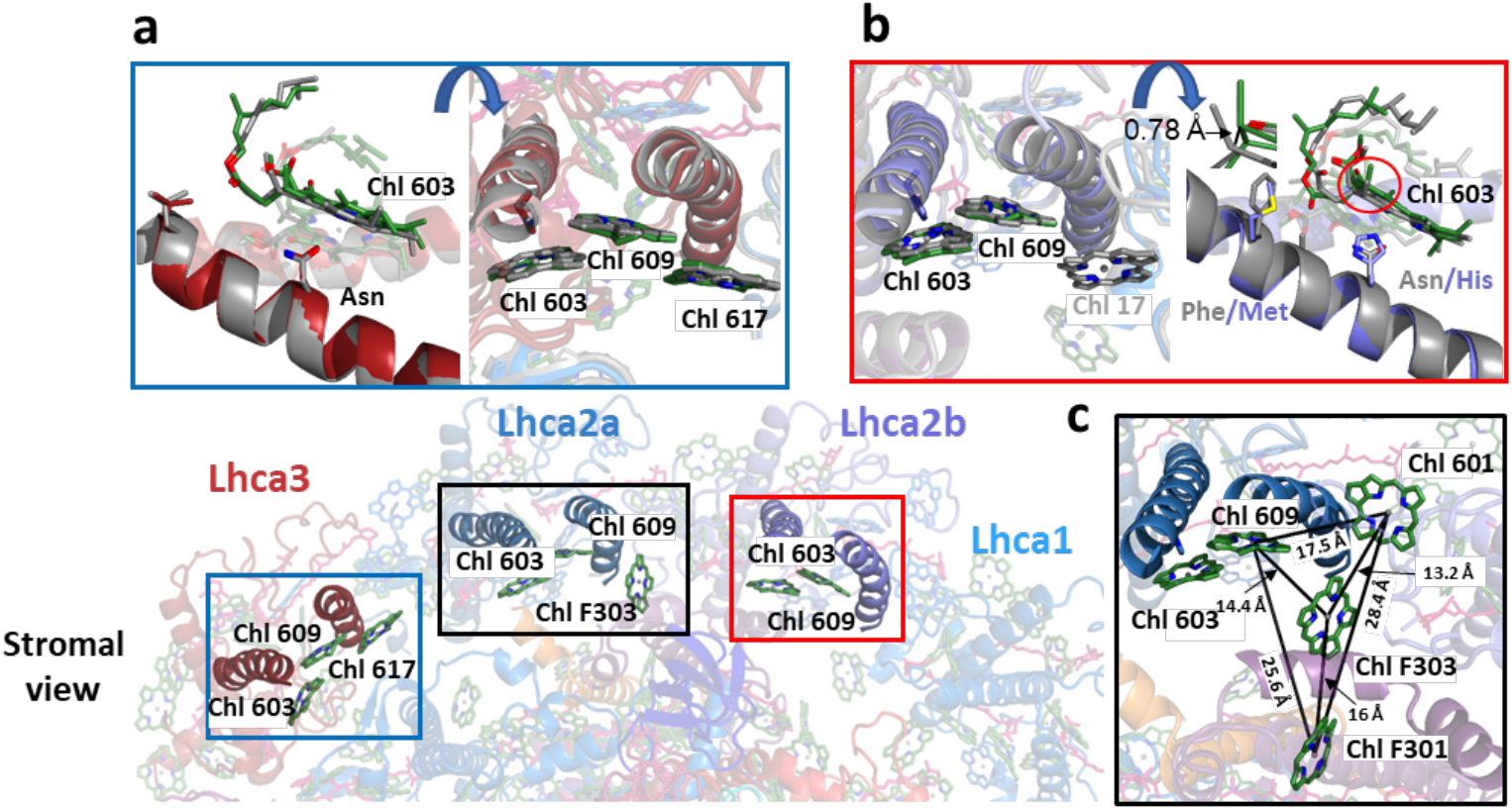
Structural differences in Chls coordination between higher plant and moss. Chl coordination is described in Lhca3 (blue square), Lhca2a (black square) and Lhca2b (red square) **a**. Superposition of the moss (red) and higher plant (gray) Lhca3 shows that Chl 603 is coordinated by Asn residue and Chl 617 is present both in the moss Lhca3 (red) and higher plant (gray). **b**. Superposition of the moss Lhca2b (purple) and the higher plant Lhca4 (PDBID:5l8r) (gray) shows that Chl 603 is coordinated by a His residue in the moss Lhca2b and by an Asn residue in the higher plant Lhca4. Chl 617 is missing in the moss Lhca2b (purple) in comparison to the higher plant (gray). **c.** Additional Chl (F303) is coordinated by the moss PsaF. Distances between subset of Chls at the interface of Lhca2a, Lhca2b and PSI are described.

An additional difference between the moss and higher plant PSI-LHCI at the interface responsible for energy transfer between LHCI and PSI is a Chl coordinated by PsaF (present in the moss PSI-LHCI), potentially contributing to energy transfer from the antenna to PSI bridging the gap between Lhca2a, Lhca2b and PSI (**figure 4c**).

### High membrane Chl *b* to Chl *a* ratio leads to increased occupancy of Chl *b* in PSI-LHCI

The absorption difference between the moss and the higher plant PSI-LHCI suggests a higher Chl *b* content in the moss complex (**figure 1d**). To eliminate the contribution of pigment protein interaction we measured the absorption of pigments extracted with 80% acetone from both complexes. The difference spectra between both extracts contains two positive peaks in the moss relative to pea around 456 nm and 645 nm (**supplementary figure 5a**) clearly indicating increased levels of Chl *b* in the moss PSI-LHCI. We analyzed the extracted pigments from both complexes using HPLC and found a small but significant difference in the Chl *b* to Chl *a* ratio between purified PSI-LHCI from moss and pea (**table 1 and supplementary figure 5b**).

**Table 1:**
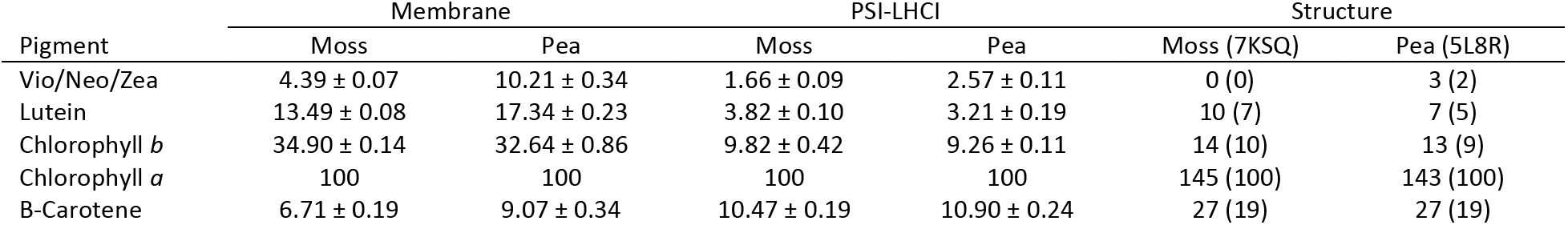
Pigment content. Pigment content per 100 chlorophyll *a* molecules and standard deviation of three biological replicas. For structures, the number in parentheses is the pigment content per 100 chlorophyll *a*.

We identified a total of three differences in Chl *b* binding in the moss PSI-LHCI structure compared to the higher plant PSI-LHCI model (**supplementary figure 6**). These differences, detailed in **figure 5a and b**, result in one net additional Chl *b* molecule in the moss PSI-LHCI structure relative to the pea (**table 1**).

**Figure 5:**
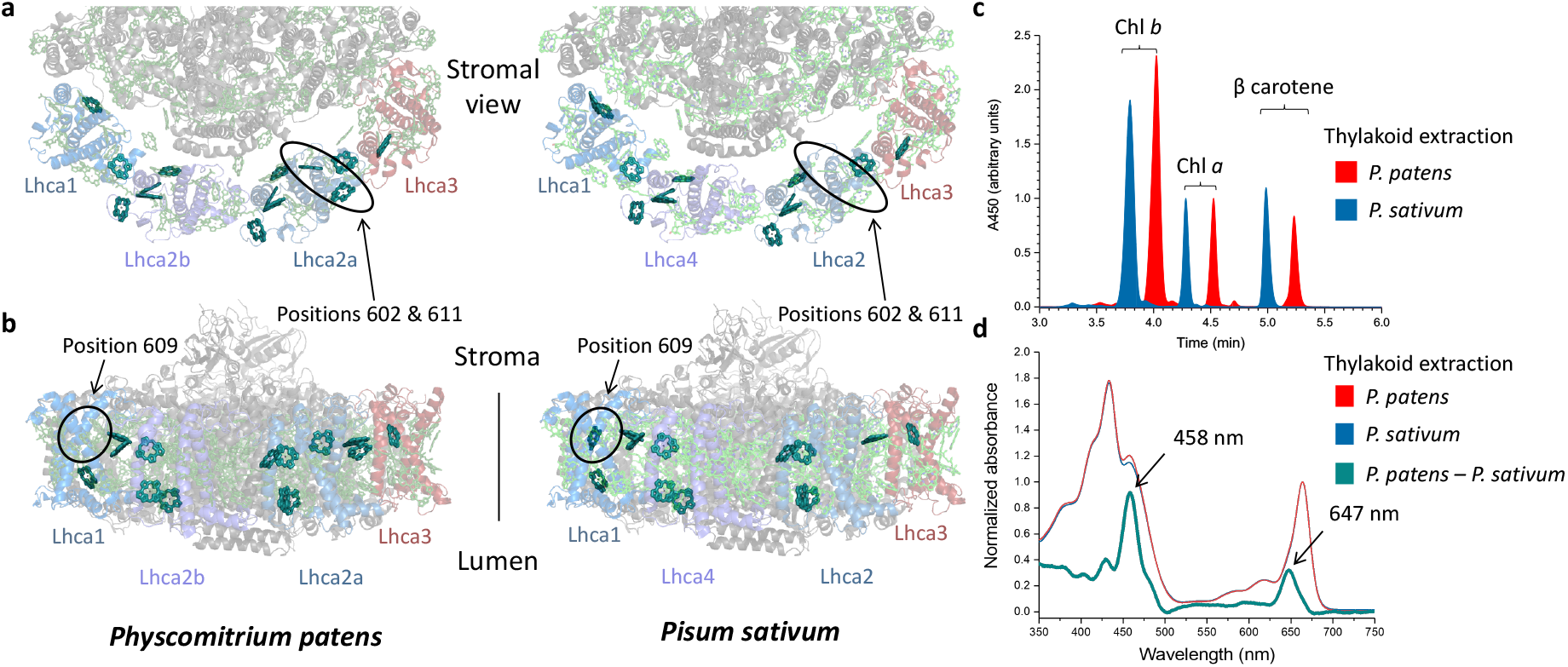
Comparison of Chl *b* positions between *P. patens* and *P. sativum* PSI-LHCI. **a**. Chl *b* molecules are shown in teal on transparent background. The moss structure contains two Chl *b* molecules in positions 602 and 611 of Lhca2a. Two chlorophyll positions that are occupied by Chl *a* in pea. **b**. Conversely, the moss structure contains a Chl *a* at position 609 in Lhca1 compared to a Chl *b* in the higher plant Lhca1. **c.** HPLC chromatogram of pigments extracted from moss and pea thylakoids. The first four minutes of the chromatograph were removed to focus on the relevant chlorophyll peaks. **d**. Absorbance spectra of pigments extracted from moss and pea thylakoid membranes.

Interestingly, no mutations that account for these Chl *b* binding differences were observed between the moss and higher plant structure. Differences in relative pigment concentrations during antenna reconstitution were shown to affect site occupancy in individual Lhca antennae^56^. In addition, the Chl *b* to Chl *a* ratio in photosynthetic organisms is known to increase when light intensity decreases^57^. We determined the Chl *b* content in both moss and higher plant thylakoid membranes (**figure 5c**). The difference spectrum of the pigments extracted from the thylakoid membranes shows two regions of higher absorption mirroring the PSI-LHCI complex difference spectrum (**figure 5d**). HPLC analysis of the extracted pigments from thylakoid membranes found a higher Chl *b* to Chl *a* ratio in the moss thylakoid membranes than in the higher plants (**table 1**). We conclude that the higher content of Chl *b* in the moss thylakoids increases the Chl *b* occupancy in the moss LHCI. It was previously shown that several sites in LHC’s can be occupied by either Chl *a* or Chl *b*^58^. Furthermore, partial occupancy explains the discrepancy between the amount of Chl *b* modeled in the moss PSI-LHCI structure as partial occupancy of a site by Chl *b* would obscure the observable differences in the Cryo-EM map.

## Discussion

The function of PSI as a Plastocyanin/Cytc6-Ferredoxin Oxidoreductase is conserved in all photosynthetic organisms, however the composition of the complex has been modified with evolution. At present, PSI structures are available from cyanobacteria, unicellular green algae, macroscopic green algae and higher plants. These structures shed light on the evolutionary forces that drive structural changes in the photosynthetic machinery. In this work we solved the structure of PSI-LHCI from the moss *P. patens*, detailing a missing link in the evolutionary line leading to higher plants.

The structure of the moss PSI is similar to the higher plant complex, as expected. Interestingly, the moss retains the PsaM gene in its chloroplast, however this subunit was not detected in the moss PSI complex previously^36^. PsaM is involved in PSI trimer stability in cyanobacteria^59^. The moss structure identified the presence of this subunit for the first time in an embryophyte PSI complex despite its monomeric state. Additional monomeric PSI that contains PsaM are observed in red algae and macroscopic green algae structures^35,51^. In those structures, PsaM seemed to mediate the binding of an additional Lhca dimer. However, in the unicellular green algae structure an Lhca dimer binds without PsaM^33^. Currently, information on the role of PsaM in eukaryotes is missing and mutagenesis studies are required to determine the importance of this subunit. The moss is probably the last member of the green lineage to contain PsaM before losing this subunit in the green lineage.

An additional feature that is retained in the moss but lost in higher plants is a short extension of the PsaF C-terminus that resembles the C-terminus found in the cyanobacteria, red algae, and green algae PSI structures. It was suggested that the moss retains an ancient version of PsaF^36^. However, the plastocyanin binding domain is highly conserved and similar to its higher plant counterpart. In addition, the moss PsaF coordinates a gap Chl at the stromal side of the interface between PSI and LHCI contributing to energy transfer (**figure 4C**).

One of the prominent differences among different PSI complexes along the green lineage is the content of red Chls in their antenna (**figure 6**)^5,21,30,48,49^. Both green algae and moss contain low energy Chls in their LHCI antenna^5,30^, however, higher plant PSI-LHCI contains the lowest energy form of Chl *a* in the Lhca4 antenna. *P. patens* appears to be adapted to low-light environments and shaded habitats. This is consistent with previous isolations of large PSI antenna supercomplexes from *P. patens* under specific growth conditions which included a red Chl containing antenna (Lhcb9) unique to moss^40,60^.

**Figure 6:**
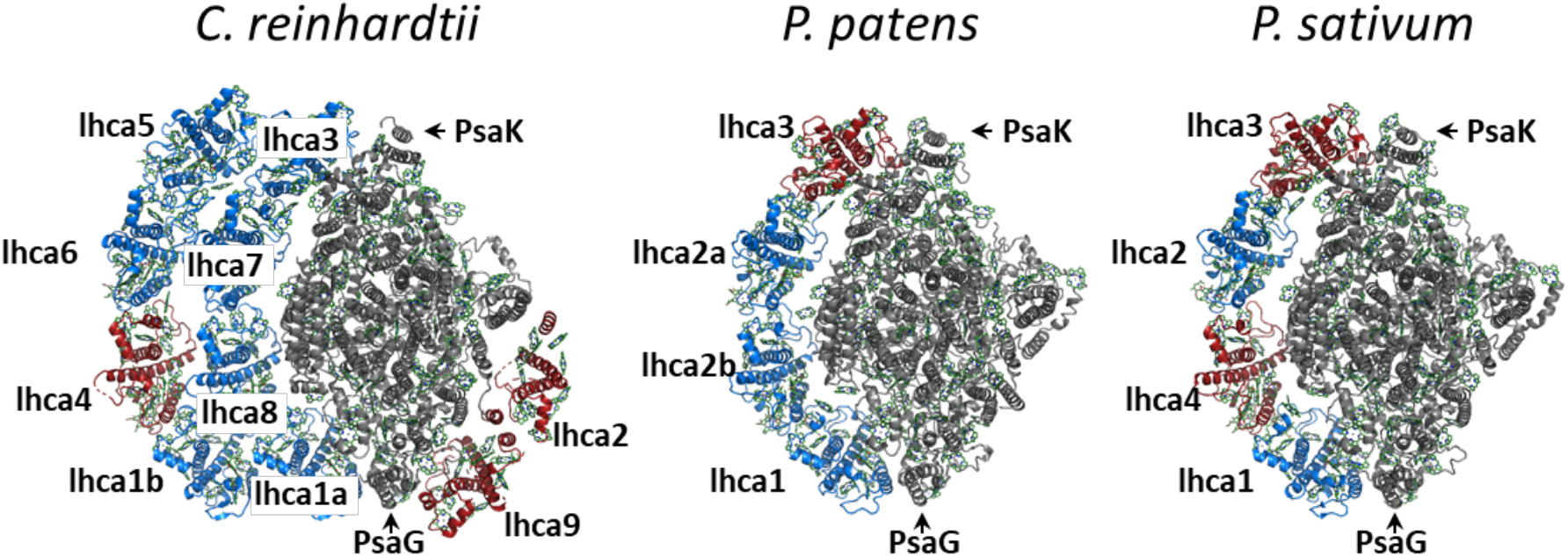
Red Chls content in the LHCI antenna in the green lineage. Lhca proteins that carry red Chls are indicated in red in the PSI-LHCI complexes from green alga (6JO5), moss (7KSQ) and higher plants (5L8R). The rest of the Lhca proteins are in blue and the core is in gray.

The composition of the core PSI-LHCI in moss has been a long standing question, and it was speculated that the moss PSI binds smaller antenna presumably composed of three Lhca proteins^5^. One subpopulation identified in our analysis contains LHCI devoid of Lhca3. The cellular role for this PSI-LHCI is not clear and its presence may be the result of the isolation procedure. However, no additional structural changes are observed in the subclass of particles, and it should be fully functional with regards to light harvesting and electron transport. The moss structure solved here revealed that a Lhca2 isoform occupies the Lhca4 protein position. As this paper was under review, a publication describing the structure of PSI-LHCI from *P.patens* at 3.2 Å resolution was published^61^. While the two structures are similar, an important difference is the assignment of the Lhca occupying the Lhca4 position. In contrast to our assignment of Lhca2b, Yan *et al*^61^ assigned Lhca5 to the same position. In supplementary figure 7, we show that the higher resolution of the map in this paper supports our assignment of Lhca2b and not Lhca5. Similar differences (albeit at lower contrast) are observed when evaluating the map published by Yan *et at*, suggesting that Lhca2b occupies this position in both structures. Lhca5 was previously implicated in the formation of the PSI-NDH supercomplex in eukaryotes^23^. It remains possible that a small subpopulation of PSI-LHCI contains Lhca5 instead of Lhca2b and contributes to the formation of PSI-NDH supercomplex.

Neither Lhca2 nor its isoform carry any red Chls, resulting in the moss’ characteristic PSI-LHCI blue shifted absorption. Intriguingly, Lhca4 is present in liverworts which are basal to mosses, suggesting that the ancestor of the moss possessed Lhca4 in the initial transition from marine to terrestrial environments and that Lhca4 loss is a late event in the evolution of *P. patens*. The contribution of red sites to antenna function under aquatic conditions, like those experienced by *P. patens*’ closely related marine ancestors, is expected to be minimal due to the absorption of far-red light by the water column^62^. At the same time, the amount of light absorbed by Chl *b* remains largely constant as blue/green light is not as readily attenuated by water. These facts can explain the higher number of Chl *b* molecules present in the moss PSI-LHCI together with the loss of Lhca4 and its associated red shifted site. We isolated PSI-LHCI from large scale, high density, liquid cultures grown under relatively low light (40 μE), light conditions favorable for Chl *b* absorption compared to previous studies which used solid media for growing cultures^40,41,61^. Pigment ratio measurements of biological replicas confirmed a higher Chl *b*/Chl *a* ratio in moss PSI-LHCI relative to pea under our experimental conditions (**table 1**). In agreement with this we assigned different Chl *b* site occupancies between the pea PSI-LHCI and the moss PSI-LHCI without any structural changes between sites (for example, additional hydrogen bonding to accommodate the formyl group in Chl *b*). This binding site plasticity was shown previously *in-vitro* for most of the pigment sites on LHC’s^63,64^. This inherent plasticity in pigment binding allows PSI-LHCI to respond to changes in the thylakoid membrane pigment pool. Consistent with this we measured a higher Chl *b* content in moss thylakoid membranes compared to pea (**figure 5, table 1 and supplementary Figure 5**). An increase in the Chl *b* to Chl *a* ratio is accepted as an adaptation to low light and shaded environments^57,65^ and has been observed in pea thylakoids^66^, however the relationship between Chl *b*/Chl *a* ratio and light intensity is species specific and not directly comparable. The organization of chlorophylls in the reaction center is highly conserved between photosynthetic organisms, in contrast to the comparatively high variability in antenna systems ^17,67^. The moss PSI structure maps light harvesting adaptations in PSI-LHCI along the green lineage from the transition to terrestrial life to current habitats.

## Data availability

The final model (PDBID 7SKQ) and map (EMD-23023) were deposited in the Protein Databank and Electron Microscopy Database, respectively. The PSI-LHCI model excluding PsaO with its corresponding map were deposited (PDBID: 7KUX, EMD-23040) as well as the model and map of PsaO alone (PDBID:7KU5, EMD-23034).

## Acknowledgements

We would like to thank Prof. Nir Ohad and Dr. Rafi Yaari for their help with moss strains, cultivation and protocols. We would like to acknowledge the use of the Titan Krios at the Erying Materials Center at Arizona State University, and the funding of this instrument by the National Science Foundation (No. MRI 1531991). This study is funded by a startup grant from Arizona State University and supported by grant number 2034021 from the National Science Foundation to Y.M.

## Methods

### Moss Thylakoid Membrane Preparation

*Physcomitrium patens* moss was grown on plates for one to two weeks under 40 μmol photons m^-2^ s^-1^ light supplied by fluorescent tubes (Agrosun 40W T12 48”) using standard BCDATG media. Plants were scraped off the plates, resuspended in BCDATG media and shredded using the VWR 200 Homogenizer on high for ten seconds. Cells were transferred to 5L bioreactors. The 5 L bioreactors were grown for one to two weeks pl shredded (VWR 200 Homogenizer on high for one minute) and transferred to 12L bioreactors. 50 L of moss culture were harvested from bioreactors, sieved, frozen in liquid nitrogen, and triturated into a powder. The powder was then suspended in STN1 buffer (30 mM Tricine-NaOH pH 8, 15 mM NaCl, 0.4 M sucrose, 1 L), sieved with cheesecloth, and centrifuged in a F9 6X1000 LEX rotor in a Sorval LYNX 6000 centrifuge (Thermo Scientific) for 20 minutes at 17,568xg. The pellet was resuspended in 200mL in 10mM Tricine-NaOH pH 8 and centrifuged in a F20-12×50 LEX rotor for 15 minutes at 41,656xg. The pellet was resuspended in 10mM Tricine-NaOH pH 8 and 150mM NaCl and centrifuged in a F20-12 × 50 LEX rotor for 10 minutes at 41,656xg. Then the pellet was resuspended in STN1 to give a chlorophyll concentration of 2.0 mg/ml and frozen at −80 C.

### Pea Thylakoid Membrane Preparation

Little Marvel pea seeds (*Pisum sativum*) from Carolina Biological Supply Company were soaked in running DI water for 1 h and planted in vermiculite trays. Peas were germinated in the dark for 5 days, then grown for 14 days in a Percival incubator (model LED41L2X) on 16/8 hour light/dark (200 μmol photons m^-2^ s^-1^) cycles at 22 °C. Leaves were harvested and approximately 150 g were ground for 20 s in a Vitamix (Low-Profile 64-ounce) with 1 L of cold STN1 buffer. The solution was filtered through cheesecloth, and the chloroplasts were pelleted by centrifugation at 1,000 g for 9 min in a Sorvall Lynx 6000 centrifuge using a Thermo Scientific F9-6×1000 LEX rotor. Chloroplasts were suspended in 500 mL of a hypotonic solution (10 mM Tricine-NaOH pH 8) and then centrifuged for 2 minutes at 500xg to remove starch. The solution was then spun at 12,000xg for 10 min to pellet thylakoid membranes. After pelleting, the membranes were resuspended in a minimal amount of cold buffer containing 0.4 M sucrose 150 mL NaCl and 10 mM Tricine-NaOH pH 8 before pelleting again at 8,000xg for 10 minutes in a Thermo Scientific F20-12×50 LEX rotor. The thylakoids were then resuspended in a minimal amount of STN2 (0.4 M sucrose, 20 mM Tricine-NaOH pH 8) and stored at −80 °C until PSI-LHCI purification.

### Moss and pea PSI-LHCI Purification

All steps were carried out on ice or in a cold room. Thylakoid membranes were solubilized by adding n-Dodecyl β-D-maltoside (DDM, Glycon) to the membranes at a 6:1 DDM to chlorophyll ratio. Insoluble material was discarded using ultracentrifugation (Ti70, 207,870xg, 30 min, 4°C). The solubilized membranes were loaded onto a diethylamino ethanol column (Toyopearl DEAE-650C) with a bed volume of 1 mL per mg of chlorophyll. The complexes were eluted using a linear NaCl gradient (15–250 mM NaCl) in 30 mM Tricine-NaOH pH 8 and 0.2% DDM. Dark green fractions were collected. The NaCl concentration was increased by 100 mM, then PEG6000 (Hampton Research) was added to a final concentration of 8% and the mixture was centrifuged in a F20-12×50 LEX rotor for 5 min at 3,214xg. The green precipitate was resuspended in 30 mM Tricine-NaOH pH 8, 50 mM NaCl with 0.05% DDM, and loaded onto a 10-40% sucrose density gradient prepared with the same buffer. Following centrifugation (Beckman SW40 rotor, 105,400xg, 18 h, 4°C), the heavier green band was collected and precipitated using 8% PEG6000 (Hampton Research) and 130mM NaCl in an Eppendorf tabletop centrifuge for 10 min at 18,407xg and 4°C. The green precipitate was resuspended in 30 mM Tricine-NaOH pH 8, 50 mM NaCl, and 0.05% DDM and loaded onto a 10– 40% sucrose density gradient prepared with the same buffer. Following centrifugation (Beckman SW60 rotor, 32,000 rpm, 18 h), the PSI-LHCI band was collected and used for subsequent experiments.

### Absorption and fluorescence spectroscopy

Absorption spectra were recorded on a Cary 4000 UV–Vis spectrophotometer (Agilent Technologies). Fluorescence spectra were recorded on a Fluoromax-4 spectrofluorometer (HORIBA Jobin-Yvon). Samples were diluted to an optical density of 0.8 and 0.1 at 680nm for absorption and fluorescence measurements respectively, using buffer containing 30 mM Tricine-NaOH pH 8, 15 mM NaCl and 0.05% DDM. The resulting spectra were normalized to the area of the chlorophyll Q bands (550-750 nm). For 77 K fluorescence measurement samples were adjusted to an OD680 of 0.1 in a buffer of 50% glycerol 30 mM Tricine pH 8.0, 15 mM NaCl, and 0.02% DDM. An Oxford instruments Cryostat was used to cool the sample to 77 K. Figures were prepared using OriginPro (OriginLab).

### Sample preparation for single-particle cryo-EM analysis

The PSI–LHCI band from the sucrose gradient was collected and precipitated using 10% PEG6000 (Hampton Research) and 120 mM NaCl. After centrifugation in an Eppendorf tabletop for 10 min at 18,407xg, 4°C the green precipitate was resuspended in 30 mM Tricine-NaOH pH 8, 50 mM NaCl with 0.05% DDM. Chlorophyll concentration of the sample was adjusted to 1.5mg chl/ml. After soaking the grid in buffer a 2-μl drop of the PSI-LHCI complex was applied to a holey carbon grid (C-flat 1.2/1.3 Cu 400-mesh grids, Protochips). The sample was vitrified by flash-plunging the grid into liquid ethane using an automated plunge freezer, a Vitrobot Mark IV (ThermoFisher/FEI) with a blotting time of 10 s. The grids were then stored in liquid nitrogen.

### Phylogenetic Analysis

The genomes of *Arabidopsis thaliana, Chlamydomonas reinhardtii, Dunaliella salina, Marchantia polymorpha, Porphyra umbilicalis, Selaginella moellendorffii, Sphagnum fallax*, and *Zea mays* were chosen to compare Lhca proteins with those in the *Physcomitrium patens* genome. All the genes annotated as Lhca from the above species were retrieved from Phytozome where they are identified by PAC numbers, (indicated in supplementary figure 4). Two moss Lhca1 genes without annotations are identified by their UniProt ID numbers. Lhca protein sequences from *Pisum sativum* and *Bryopsis corticulans* were retrieved from the PDB (PDB ID 5L8R and 6IGZ, respectively^1,2^). The genes were aligned using Muscle^3^ with default settings using Jalview^4^. The tree was constructed in MegaX using the Maximum Likelihood method with default settings^5^.

### Pigment analysis

The HPLC protocol was taken from De Las Rivas *et al*.^6^. Pigments from thylakoids membranes and purified PSI-LHCI samples from moss and pea were extracted into 80% acetone by combining sample and 100% acetone at a 1:4 ratio. Extracted samples were incubated at room temperature for 5 minutes, and then centrifuged at 21,130 xg for 10 minutes at room temperature in an Eppendorf 5424 R benchtop centrifuge. The organic top layer was removed and filtered using a Millex-GV 0.2 μm 4 mm filter. Pigments were separated using an Agilent ZORBAX Eclipse Plus C18 column (4.6 × 100 mm, 3.5 μm) calibrated to 20 °C on an Agilent Technologies 1100 series HPLC fitted with a G1322A degasser, G1311A quaternary pump, G1313A ALS autosampler, G1314A VWD detector, and G1364A AFC automatic fraction collector. The sample injection volume was 60 μl, detection wavelength was set to 450 nm, and flow rate was 2 mL/min. The column was equilibrated in mobile phase A (82:18 Acetone:H_2_O) prior to Injection. Mobile phase A was used for the first two minutes of run, followed by mobile phase B (7:0.96:0.04:2 Acetonitrile:MeOH:H_2_O:Ethyl acetate) which was pumped for 1 minute, followed by mobile phase C (7:0.96:0.04:8 Acetonitrile:MeOH:H_2_O:Ethyl acetate) which was pumped until the end of run at 6 minutes post injection.

### Weighted residue distances

To calculate distances between terminal nitrogen and oxygen atoms in adjacent residues in figure 3b and c we considered all possible rotamer assignments compatible with the experimental map. Each rotamer pair was refined into the map in coot^7^ before distance measurement. Finally, the weighted average distance was calculated together with the standard deviation.

### Cryo-EM data acquisition

The cryo-EM specimens were imaged on a Titan Krios transmission electron microscope (ThermoFisher/FEI). Electron images were recorded using a K2 Summit direct electron detect camera (Gatan) at super-resolution counting mode. The defocus was set to vary from −1 to −3 μm. The nominal magnification was ×47,600, corresponding to a super-resolution pixel size of 0.52 Å at the specimen level. The counting rate was adjusted to 8 e^-^/Å^2^ s. Total exposure time was 8 s, accumulating to a dose of 64 e^-^/Å^2^.

### Data processing

A flowchart describing data handling is shown in Supplementary figure 1. MotionCor2^8^ was used to register the translation of each sub-frame, and the generated averages were rescaled to a 1.04 Å/px (2X binning) and dose-weighted. Contrast transfer function (ctf) parameters for each movie were determined using CTFFIND4^9^. Relion3.1 was then utilized for subsequent data processing^10^. A set of manually picked particles (~1,000) was subjected to several rounds of unsupervised 2D classification. Six class averages representing different orientations of the expected particle were selected and used as templates for the automated particle selection procedure as implemented in Relion which yielded 456,923 particles. This particle set was subjected to several rounds of unsupervised 2D classification (Relion), leading to a set of 267,486 particles which were extracted from the original micrographs as boxes of 280 pixels at 1.04 Å/Px. This particle set was reconstructed to an initial resolution of 3.5 Å. The dataset was further filtered using four repeated rounds of 2D classification without angular refinement followed by ctf refinement and 3D reconstruction bringing the reconstruction resolution to 3.24 Å. Following Bayesian polishing in Relion^11^, two more rounds of per particle ctf refinement^12^ followed by 2D classification without refinement and 3D reconstruction led to 127,092 particles and a 2.8 Å resolution reconstruction. FSC curves and the final sharpened map (using a b factor of −80) were obtained from the post processing step in relion. Local resolution was estimated using ResMap^13^. The distribution of different views in the final reconstruction was measured using cryoEF^14^.

### 3D classification and map generation

3D classification of the dataset identified one major class containing 114,608 particles. This particle set was used to reconstruct the final PSI-LHCI map (EMD-23040) and used for focused classification. Focused classification using a mask surrounding LHCI identified ~9% of the particles which lacked Lhca3, reconstructions omitting this class were indistinguishable from ones obtained using the full dataset. Focused classification using a spherical mask positioned around the PsaL end revealed four significant classes in the dataset described in supplementary figure 2d. Class number 6 contained clear additional densities for PsaO and an additional, inverted, Lhc protein. Other classes were mainly differentiated in the density of the PsaH subunit which was missing in about 10% of the particles.

### Modeling

The plant PSI-LHCI (PDBID 5L8R) model^2^ was docked into the map using Phenix. The sequence was manually changed to the corresponding *P. patens* subunits by manually adjusting the model in coot^7^. Assignment of Lhca2a and Lhca2b is described in the main text. The model was refined using Phenix^15^. For PsaO modeling, the map from class6 in supplementary figure 2d (EMD-23034) was used to dock an initial PsaO model taken from the *D. salina* PSI-LHCI structure^16^ (PDBID: 6SL5). After rigid body docking done in chimera^17^ the sequence was changed to the *P. patens* sequence and the model was manually fitted to the map using coot^7^. For side chain modeling the map was dynamically sharpened in coot using a b-factor range of −90 to −140. Finally, the map was exported from coot (sharpened with a b factor of −120) and combined with the PSI-LHCI map using the phenix.combined_focused_maps tool together with a combined PSI-LHCI-PsaO model. This map was masked in Chimera to remove the detergent belt noise and the additional Lhc density near PsaO, and then deposited in the PDB and EMDB (PDBID: 7SKQ, EMD-23023).

