## Supplemental material for "The structure of the *Physcomitrium Patens* Photosystem I Reveals a Unique Lhca2 Paralogue replacing Lhca4"

### Supplementary Figure 1

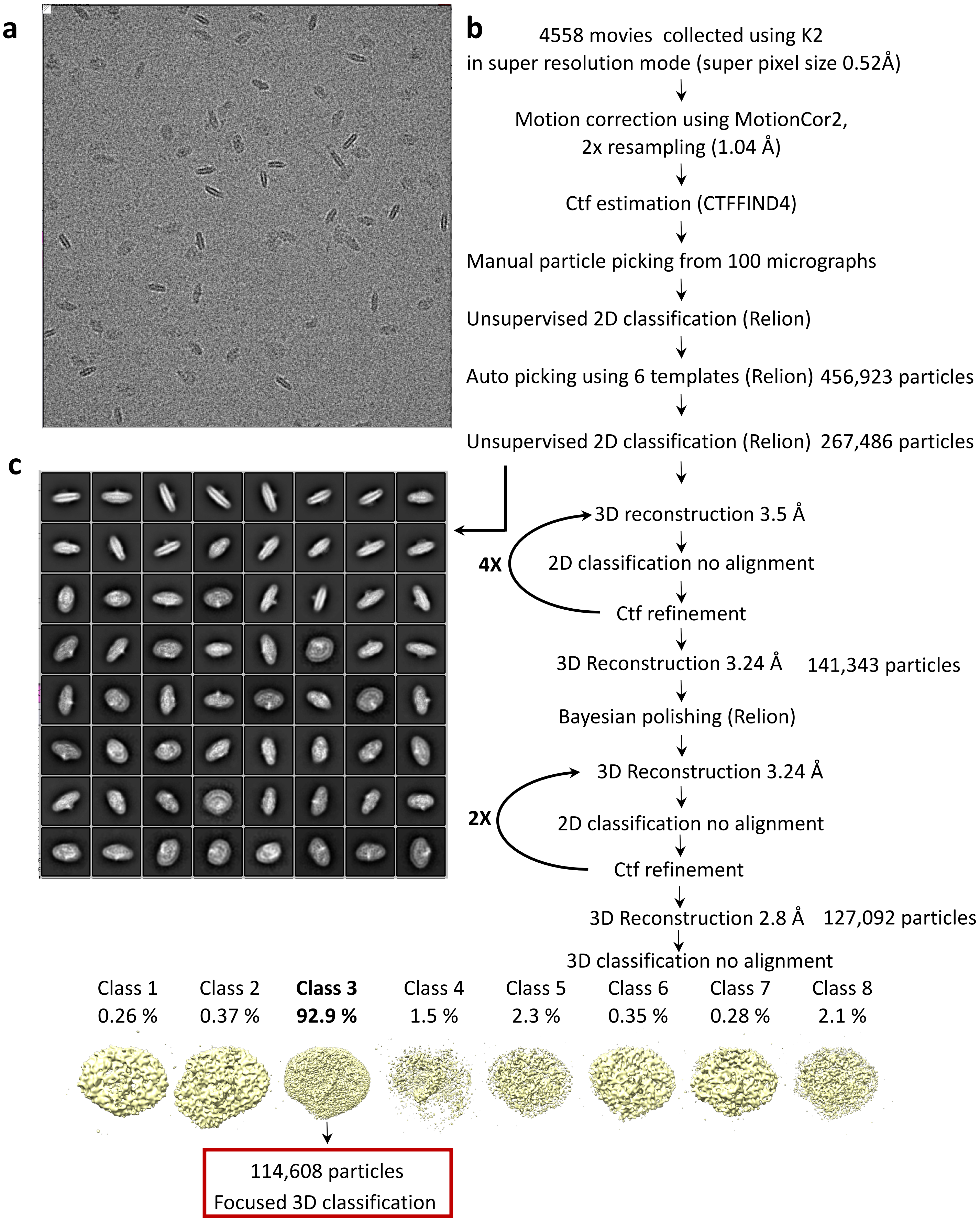


**Supplementary figure 1. Image quality and processing strategy.**

**a**. A representative micrograph. PSI-LHCI particles are visible in different orientations as projections in vitreous ice on the grid. **b**. Data collection and image processing strategy before focused classification, further details can be found in the material and method section. The final particle dataset highlighted in the red square. **c**. 2D class averages from the indicated processing stage.

Supplementary Figure 2

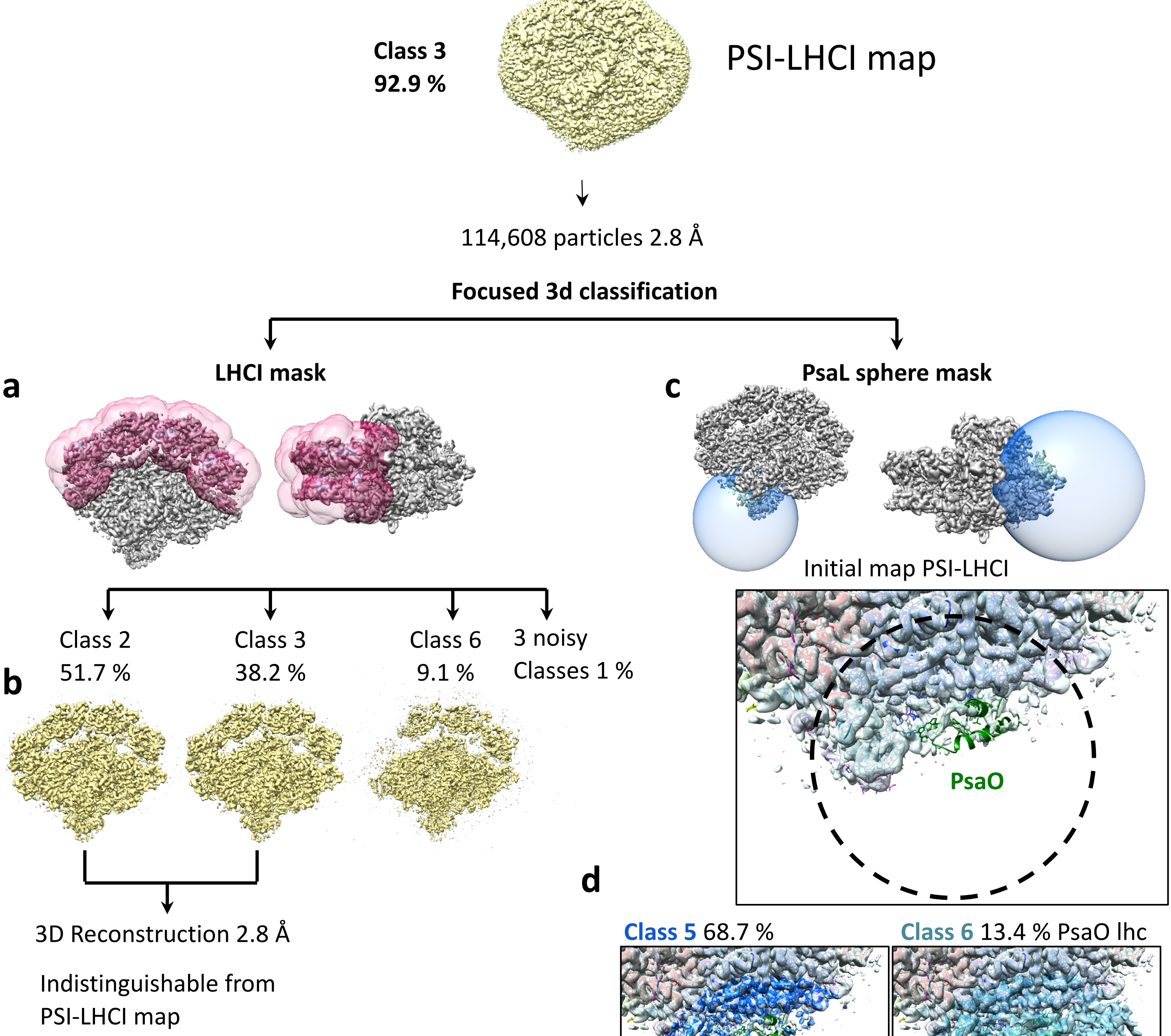


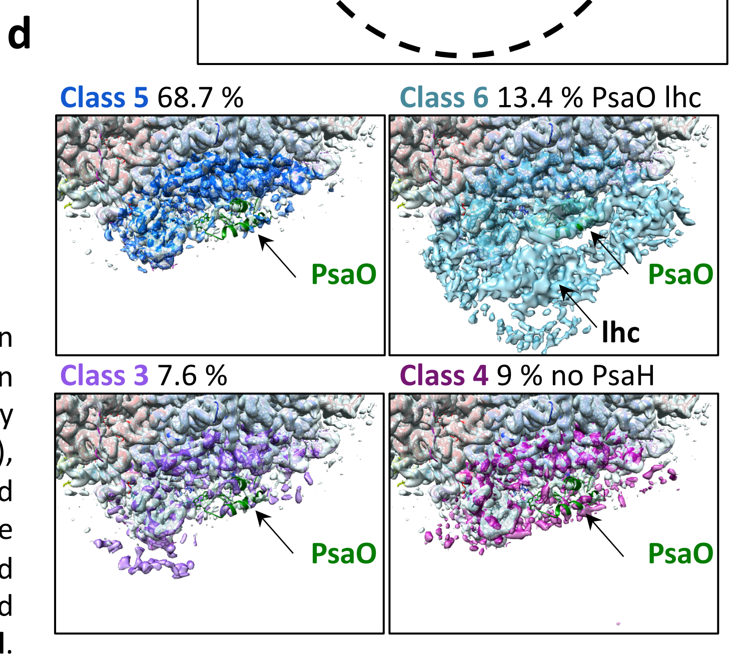


**Supplementary Figure 2:** Focused classification results. **a**. The mask used for focused classification on LHCI shown in red. **b**. We used six classes to classify the particle set in place (no orientation refinement), the three significant classes are shown. Classes 2 and 3 were used to generate a map which refined to the same resolution as the map before focused classification. **c**. The spherical mask used for focused classification of the PsaL end of PSI-LHCI. **d**. classification of the dataset resulted in four major classes. One (class 6) containing PsaO (protein model in green, map in blue, original PSI-LHCI map in light grey) and an inverted, truncated, Lhcb1 (see supplemental figure 3 for assignment). A second class (class 4) lacked the PsaH subunit. The two remaining classes (class 5 and 3) contained comparable densities in all the assigned subunits.

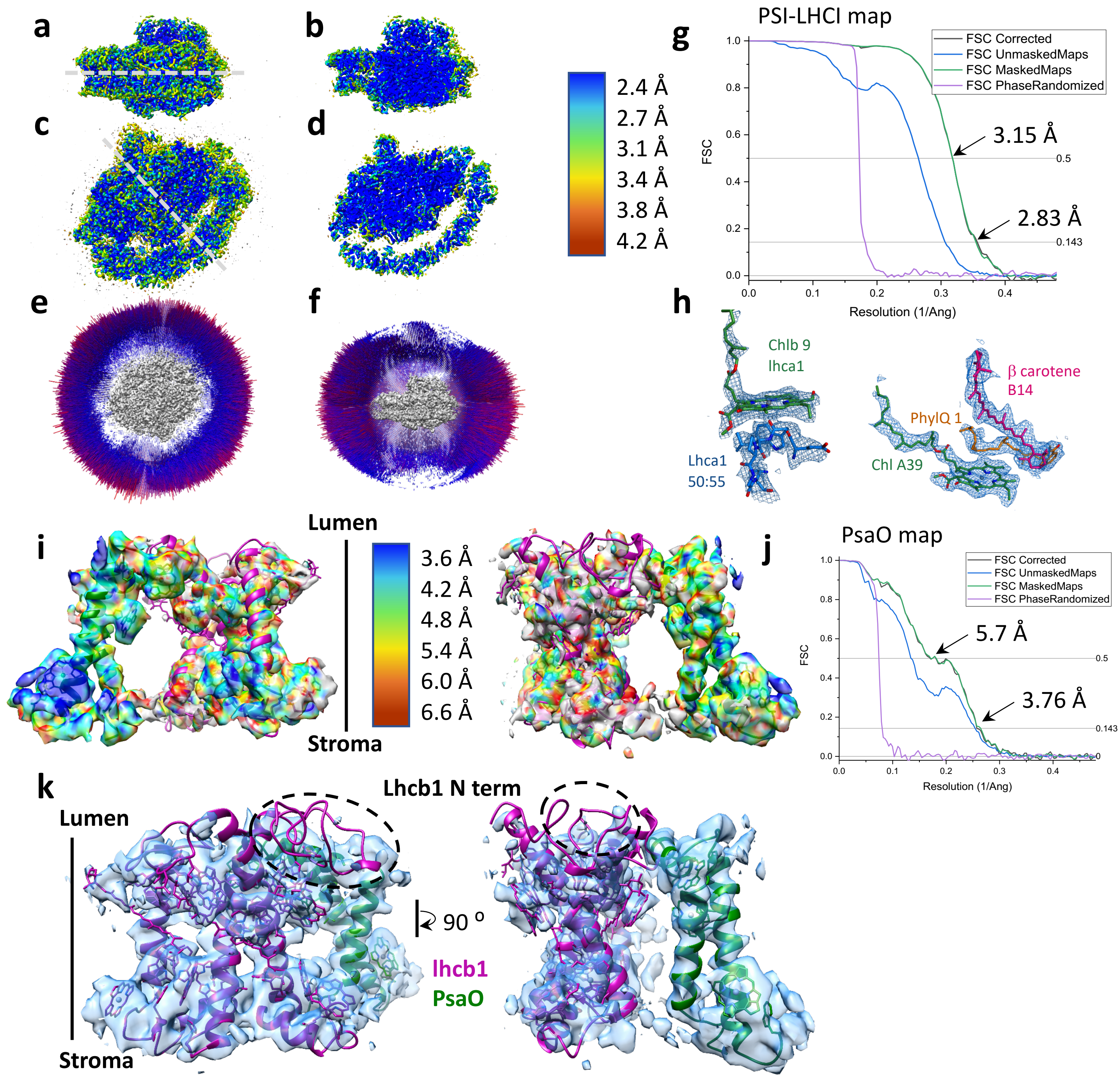


**Supplementary figure 3:** Resolution estimates and map examples. **a** and **b**. Side views of the local resolution map (with 'd' showing a section through the map, as indicated by the dashed line in 'a'). **c** and **d**. The local resolution map viewed from the stromal side ('c') (with 'b' showing a section through the map, as indicated by the dashed line in 'c'). **e** and **f**. Distribution of views in the final reconstruction (all 114,608 particles). While some views from above the membrane plan were less frequent or missing, the overall efficiency calculated using cryoEF was still 0.72. **g**. Fourier shell correlations between half maps reconstructed from the final particle set (114,608 particles). **h**. Map examples from the PSI core (left) and LHCI (right). **i**. Local resolution map of PsaO and the unidentified lhc obtained from class 6 (Supplementary figure 2d). **j**. Fourier shell correlations between half maps in the class 6 reconstruction. **k.** Unsharpened maps around PsaO (in green) and the truncated Lhcb1 (in purple) obtained from class 6. The model for lhcb1 is a monomer from PDBID: 2BHW and was refined into the map as a rigid body. Map around the N terminus and the first turn of helix 1 is clearly missing. The orientation of this Lhcb1 in the membrane is inverted as is clearly determined from the map model fit of three remaining helices and pigment clusters.

### Supplementary Figure 4

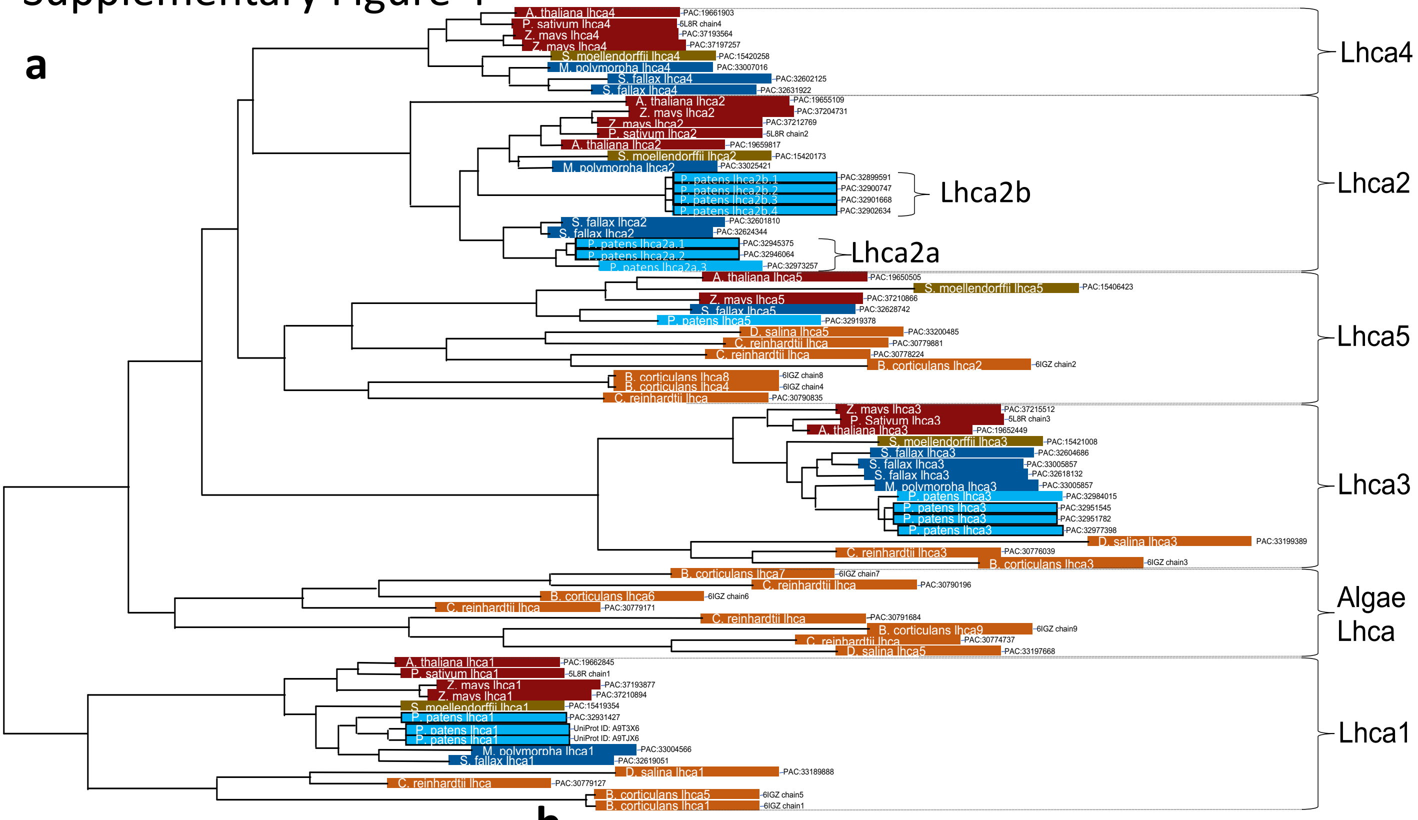


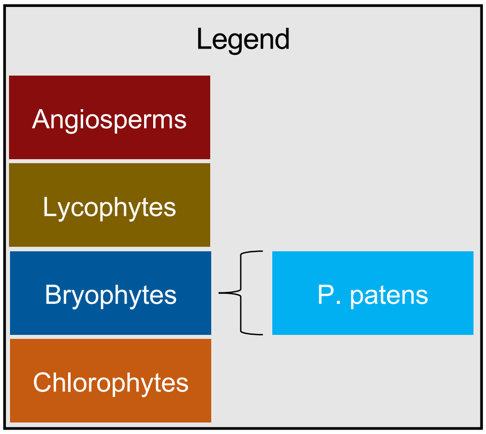


b

| Lhca name | PAC Number |
|---|---|
| P. patens lhca2b.1 | 32899591 |
| P. patens lhca2b.2 | 32900747 |
| P. patens lhca2b.3 | 32901668 |
| P. patens lhca2b.4 | 32902634 |
| P. patens lhca2a.1 | 32945375 |
| P. patens lhca2a.2 | 32946064 |
| P. patens lhca2a.3 | 32973257 |

c

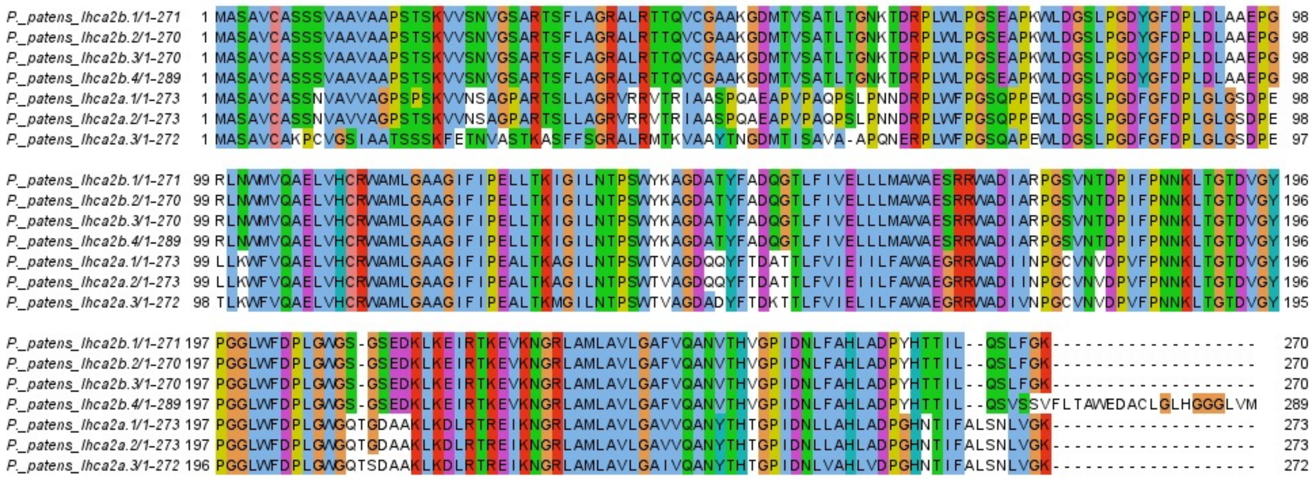


**Supplementary figure 4. Phylogenetic tree of Lhca genes.**

**a.** Phylogenetic tree. **b**. *P. patens* has seven Lhca2 genes which cluster into two families, Lhca2a and Lhca2b. Lhca4 genes present in other Bryophytes (but missing in *P. Patens*) are shown in the tree for *M. Polymorpha* and *S. fallax*. (b) Table of Lhca names and searchable PAC numbers. **c.** Alignment of all moss Lhca2 genes.

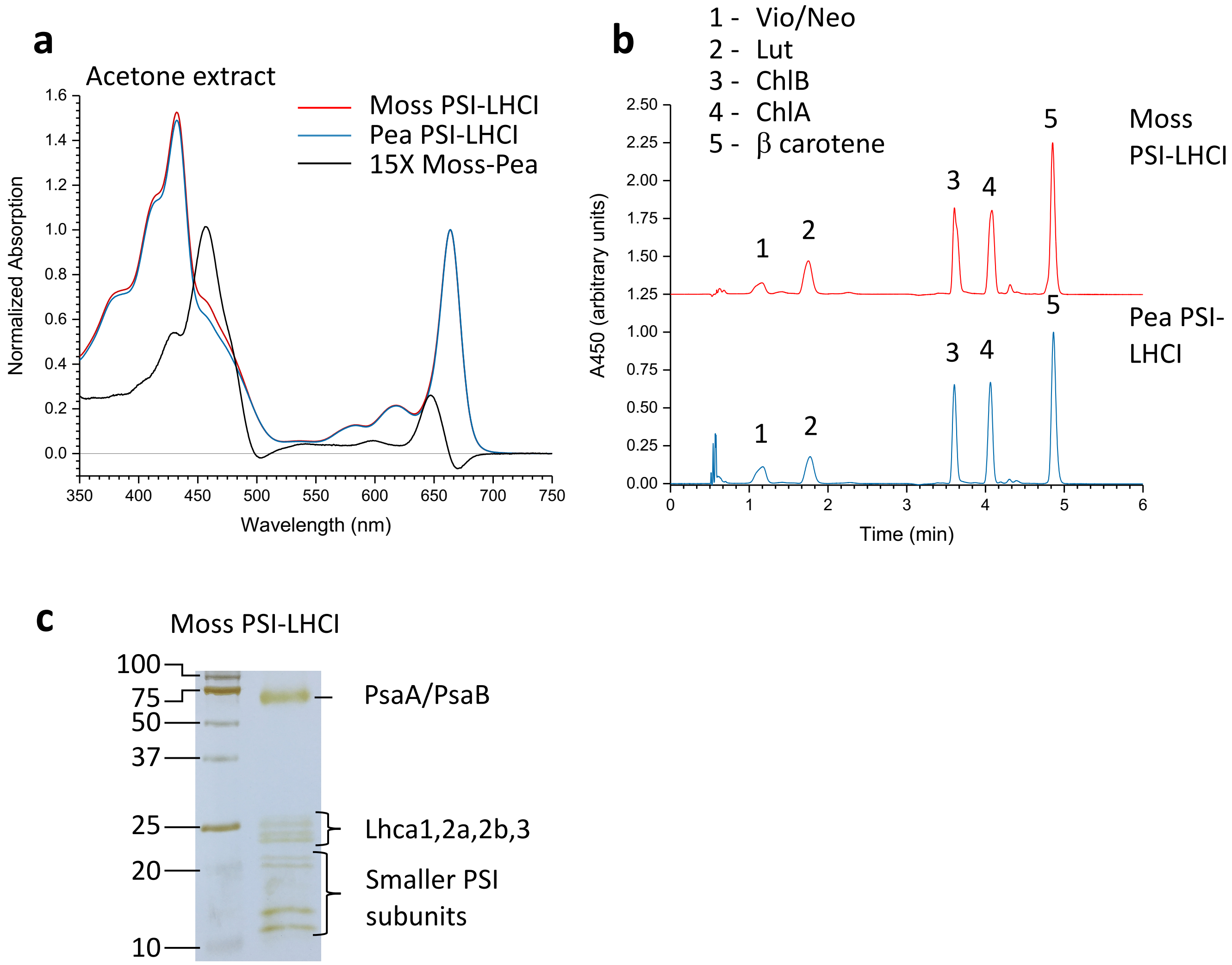


**Supplementary figure 5:** Pigment composition. **a**. Absorption spectra of pigments extracted from PSI-LHCI from moss and plants in 80% acetone together with their difference spectra in light blue (multiplied by 15). The difference spectra indicate an increase in the Chl b to Chl a ratio in the moss PSI-LHCI. **b**. Averaged (3 biological replicas) chromatogram of pigments extracted in 80% acetone and separated using HPLC. **c**. SDS-PAGE of *P. patens* PSI-LHCI preparation.

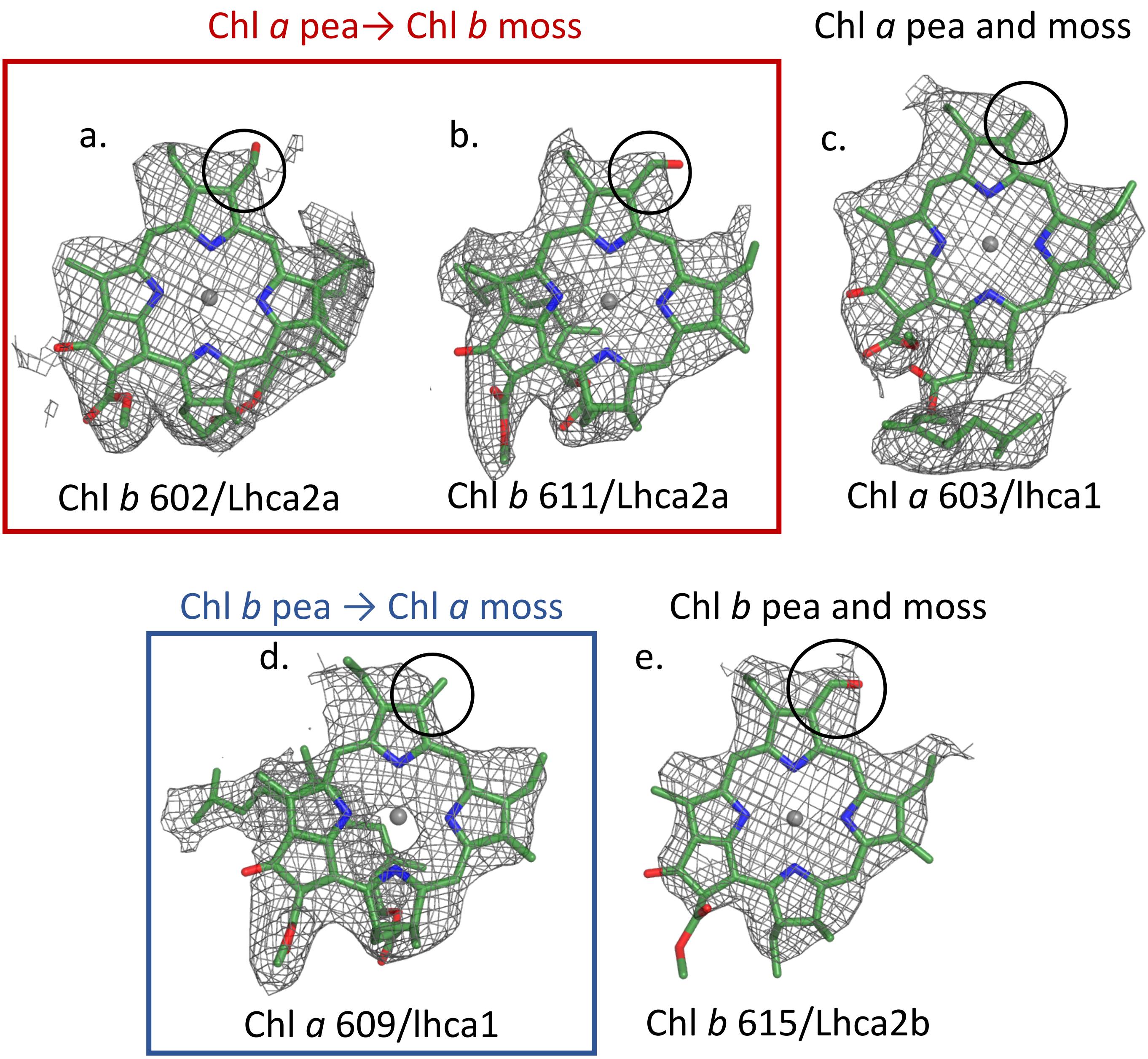


**Supplementary figure 6. Chl *b* assignment. a.** and **b.** Chl *b*'s assigned in the *P. patens* Lhca2a which are assigned as chl *a* in *P. sativum*. Map features around the chl *b* formyl group are circled. **c.** A comparable chl *a* map sample (Lhca1 Chl 603, which is assigned as chl *a* in both *P. patens* and *P. sativum)* showing the same map region circled. **d.** Map density around a plant chl *b* site (Lhca1 Chl 609) which was assigned as chl *a* in the *P. patens* PSI-LHCI (while the density suggest that a pheophytin may occupy this position, no evidence for pheophytin was detected in HPLC measurements). **e.** Map density around a chl *b* site shared between both *P. patens* and *P. sativum*.

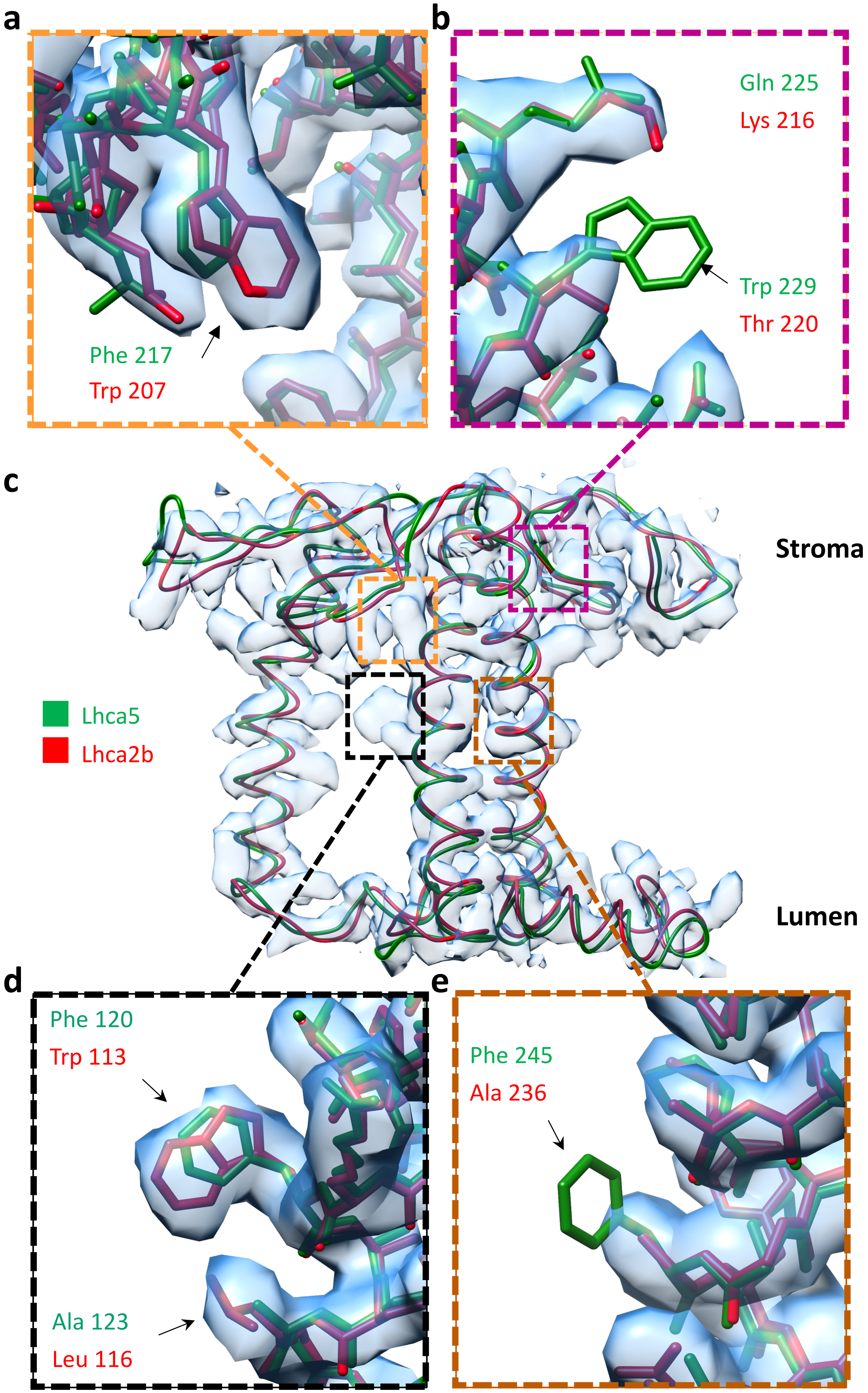


**Supplementary figure 7. Lhca2b model-map fit is preferable to Lhca5 on multiple positions.** Lhca5 was superimposed on Lhca2b in the experimental 2.8 Å map for comparison. The map is shown as a semitransparent surface using Chimera (map level 1.91). **a, b, e,** and **d**. Detailed views of residues in boxed regions of the superimposed structures that illustrate a better map to model fit using Lhca2b sequence compared to Lhca5. **c.** Comparing the backbone configuration of the two Lhc's reveals the overall configuration common to all Lhc's.
